# SITH: an R package for visualizing and analyzing a spatial model of intratumor heterogeneity

**DOI:** 10.1101/2020.07.10.198051

**Authors:** Phillip B. Nicol, Dániel L. Barabási, Amir Asiaee, Kevin R. Coombes

## Abstract

**Motivation:** Cancer progression, including the development of intratumor heterogeneity, is inherently a spatial process. Mathematical models of tumor evolution can provide insights into patterns of heterogeneity that can emerge in the presence of spatial growth.

**Summary:** We develop SITH, an R package that implements a lattice-based stochastic model of tumor growth and mutation. SITH provides 3D interactive visualizations of the simulated tumor and highlights heavily mutated regions. SITH can produce synthetic bulk and single-cell sequencing data sets by sampling from the tumor. The streamlined API will make SITH a useful tool for investigating the relationship between spatial growth and intratumor heterogeneity.

**Availability and Implementation:** SITH is a part of CRAN and can thus be installed by running `install.packages(“SITH”)` from the R console. See https://CRAN.R-project.org/package=SITH for the user manual and package vignette.

## 1. Introduction

A comprehensive understanding of how intratumor heterogeneity (ITH) develops is critical for effective cancer diagnosis and treatment (Stanta and Bonin, 2018). Mathematical models of cancer evolution are a promising approach for studying ITH and are free of the ethical and logistical questions associated with collecting clinical data (Beerenwinkel et al., 2015). Although the general evolutionary dynamics of cancer growth are well-characterized (Michor et al., 2004), little is known about the effect of spatial growth on ITH. Developing an in-silico model that captures the evolution of a spatially embedded tumor would be a starting point for investigating this relationship. Such a model may also be useful for developing novel statistical methods which can account for samples collected from a spatially heterogeneous tumor.

Our package ‘A Spatial model of Intra-Tumor Heterogeneity (SITH)’ implements a stochastic model of 3D tumor growth and mutation. The growth model is inspired by Waclaw et al. (2015) and similar models have recently been applied to study the limitations of sequencing data in providing a representative sample of a spatially heterogeneous tumor (Chkhaidze et al., 2019; Opasic et al., 2019). SITH allows users to simulate tumors with millions of cells in under a minute and provides useful features for analyzing the results. SITH can also produce synthetic single-cell and bulk sequencing data sets from the simulated tumor. SITH may prove useful in uncovering spatial biases in statistical methods, or as a basis for improving sampling techniques to ensure that a representative subset of the tumor population is obtained.

## 2. Features

The core function of SITH is `simulateTumor()`, which implements a stochastic model of tumor growth and mutation where cells occupy sites on a 3D lattice. The spatial component limits cell replications to unoccupied adjacent sites on the lattice. During the replication process, the daughter cells may acquire neutral or advantageous genetic alterations. The user can specify cell replication rate, death rate, mutation rate, and selective advantage conferred to driver mutations. See *Supplementary Information* for more details on the model as well as the simulation algorithm used.

### 2.1. Visualization of the simulated tumor

In-silico tumors produced by SITH can be rendered in an interactive 3D environment through the rgl package (Adler and Murdoch, 2020). As shown in Figure 1A, we have implemented two modes to visualize the tumor. On the left, each unique genotype is assigned a distinct color. On the right, cells are colored by their mutational burden, with blue corresponding to few and red corresponding to many mutations. To look inside the tumor, `plotSlice()` allows the user to view any 2D cross-section.

**Figure 1:**
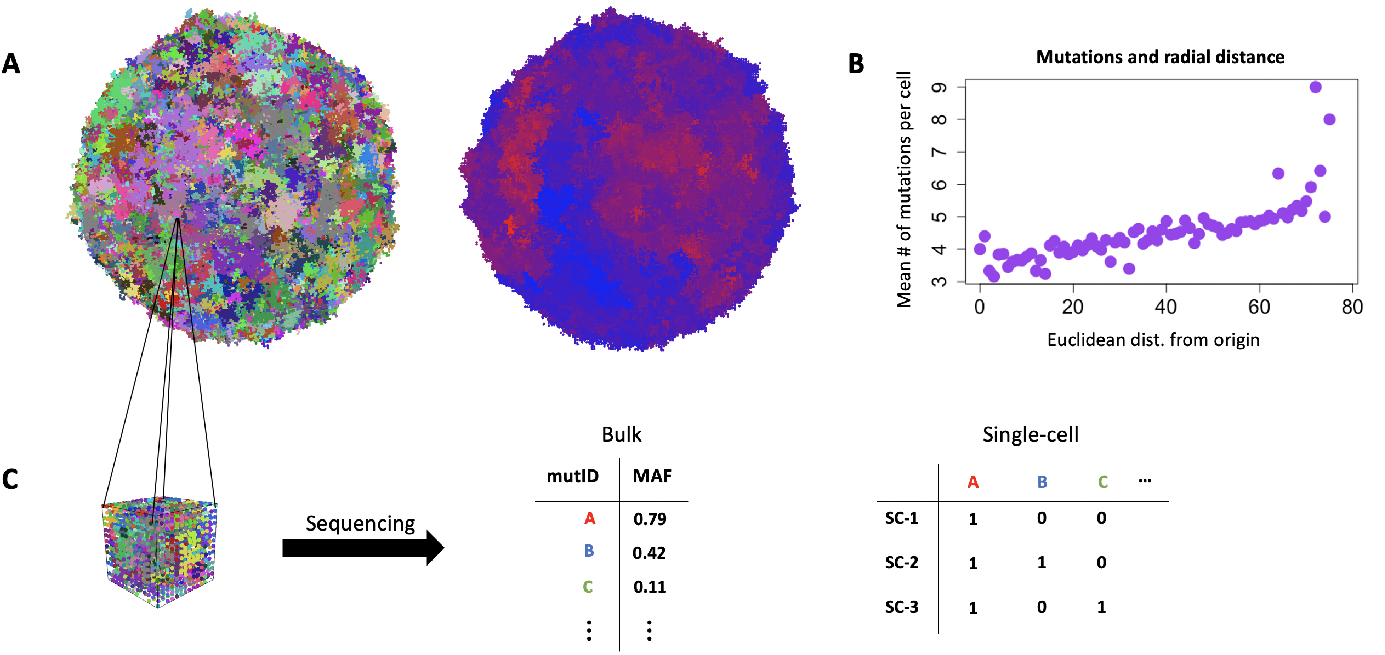
The main features of SITH. **A**: 3D snapshots of a simulated tumor (10^6^ cells). On the left, each unique genotype is assigned a color. On the right, regions with high mutation are colored red while regions with low mutation are colored blue. **B**: A plot of average mutations per cell as a function of Euclidean distance from the origin. **C**: A cube is selected from the tumor and sequenced, returning bulk or single-cell data.

### 2.2. Quantifying the spatial distribution of mutants

A crucial unknown for sampling tumors is how spatial growth biases the system’s distribution of genetic diversity. SITH was designed to provide a sandbox for asking questions about the spatial distribution of mutants within a tumor. `spatialDistribution()` can produce relevant measurements of spatial heterogeneity, which can be either plotted through SITH or output as data for further study. The function catalogues the average number of mutations per cell at varying radial distances, as plotted in Figure 1B. The plot suggests that highly mutated cells are more commonly found near the boundary of the tumor. Another included measure of heterogeneity compares the average Jaccard similarity of cells separated by varying distances. One might expect that cells in the same neighborhood share more genetic similarity than cells on opposite sides of the tumor.

### 2.3. Synthetic sequencing data

Bulk sampling is modeled as selecting an *n* × *n* × *n* cube from the tumor to be sequenced (Figure 1C), which returns mutation allele frequencies (MAF) for each mutation. Note that unoccupied lattice sites are assumed to be normal tissue, and thus the MAF may be less than 1 even if a mutation is clonal. This procedure is clinically realistic, since it is oftentimes difficult to deconvolve cancer cells from normal tissue (Opasic et al., 2019). `bulkSample()` makes multiregion bulk sampling easy by randomly selecting cubes or by allowing the user to input cube location. To simulate fine needle aspiration, `randomNeedles()` sequences random 1D cross sections of the tumor.

With `singleCell()`, the user can create synthetic single-cell sequencing data sets by either selecting cells randomly or at specified positions. Due to artifacts of sequencing technology, single-cell data sets are expected to have high noise rates (Zafar et al., 2018). To account for this, `singleCell()` allows the user to introduce false negatives and positives at a specified rate.

## 3. Discussion

With a straightforward API that can be used entirely within R, SITH provides a biologically motivated simulation of spatial tumor growth, coupled with methods for measuring ITH. Synthetic data generated from SITH can serve as the ground truth for benchmarking various computational methods. For example, the single-cell data could be used as input to various phylogenetic tree reconstruction algorithms, such as those presented in Schwartz and Schäffer (2017). Similarly, SITH can be used to test the accuracy of algorithms designed to estimate subclonal composition, since the true MAF for each mutation is provided.

Planned extensions of SITH include simulations of metastatic seeding and treatment. By analyzing cells near the tumor periphery, SITH can provide insights into the likely genetic compositions of metastases. Incorporating simulations of treatment will allow for comparisons of the cancer recurrence time under a variety of surgical and therapeutic procedures.

## Acknowledgements

D.L.B. was supported by NIH NIGMS T32 GM008313.

