## Supplementary Information for "SITH: an R package for visualizing and analyzing a spatial model of intratumor heterogeneity"

### 1 Model details

We model tumor growth as a multi-type birth-death process where cells occupy sites on the three-dimensional integer lattice. Each cell is given a birth rate $b$ and death rate $d$ such that the time until cell replication or cell death is exponentially distributed with parameters $b$ and $d$, respectively. A cell can replicate if at least one of the six sites adjacent to it is unoccupied. Note that the replication rate does not depend on the number of free neighbors. Each time a cell replication occurs, both daughter cells acquire $Pois(u)$ genetic alterations. $u$ is usually small and was estimated to be $\approx 0.01$ by Bozic et al. (2010). We follow the "infinite alleles model" (Kimura and Crow, 1964) in assuming that each alteration occurs only once, so that each mutation event introduces a unique allele in the population.

An alteration is a driver mutation with probability $u_d$ (corresponding to parameter `du` in the package) and is a passenger mutation otherwise. To model the selective advantage conferred to driver mutations, we assume that the birth rate of a cell depends on the number of driver mutations present in its genotype. A cell with $k$ driver mutations is given birth rate $bs^k$ where $s > 1$ and $b$ is the initial, or wild-type, birth rate. As noted by Waclaw et al. (2015), this is equivalent to decreasing $d$ as the dynamics of the model are determined by $b/d$.

Since cancerous tumors are believed to often begin with a single mutated cell Cooper (2000), the initial state of our model is a single cell at the origin with $b > d$.

### 2 Simulation algorithm

We simulate our algorithm using the Gillespie algorithm (Gillespie, 1977). Suppose there is a population of $n$ cells at time $t$. For cell $i$, the time until the next event, whether that be death or replication, is distributed as $Expo\,(b_i + d_i)$.[1] Since the waiting times for the next event in each cell are independent, the

[1] We assume throughout that although a cell cannot replicate if it has no available neighbors, it may "attempt" to replicate even if no neighbors are free.

time until the next event is distributed as $Expo\left(\sum_{i=1}^{n} b_i + d_i\right)$. The probability that the next event involves cell $i$ is $(b_i + d_i)/\sum_{i=1}^{n}(b_i + d_i)$, by competing exponentials. These observations inspire the following procedure to simulate the model:

1. Initialize a cell with $b > d$ at the origin.

2. While the total population of cells $n$ is less than $N$, select cell $i \in [n]$ with probability $(b_i + d_i)/\sum_{i=1}^{n}(b_i + d_i)$. With probability $b_i/(b_i + d_i)$ cell $i$ will attempt to replicate. Otherwise, cell $i$ will die and we proceed to step 4.

3. Cell $i$ is selected to replicate. If none of the 6 adjacent lattice sites are unoccupied, then proceed to the next step. Otherwise, there is at least one adjacent site that is unoccupied. Randomly select one of these sites. Place a daughter cell with the same genotype as the parent at the selected site. Next, both the parent and the daughter are independently given $Pois(u)$ additional genetic alterations. If any of these are driver mutations, then update the birth rate accordingly.

4. Update the time as $t + X$, where $X \sim Expo\,(np_{\max})$, where $p_{\max} = \max_{i\in[n]}(b_i + d_i)$.

To implement step 2, we must select a cell from the population according to the desired probability distribution. The standard approach to do this is by using inverse transform sampling. This requires computing the cumulative sum over a vector of length $N$, which will be costly as $N$ becomes large. For realistic parameter values (i.e., $b \approx d$, $s \approx 1$ (Bozic et al., 2010)), we expect the distribution to be "approximately uniform." For this reason, we use rejection sampling to perform step 2. For each iteration of rejection sampling, a random cell $i$ is selected uniformly from the population. Next, we obtain a sample $u$ from the uniform distribution over $[0, p_{\max}]$, where $p_{\max}$ is as above. If $u < b_i + d_i$, then cell $i$ is selected to replicate or die. Otherwise, we proceed to the next iteration of rejection sampling.
